# Interferometric Image Scanning Microscopy for label-free imaging at 120 nm resolution inside live cells

**DOI:** 10.1101/2025.09.19.677416

**Authors:** Michelle Küppers, W. E. Moerner

## Abstract

Light microscopy remains indispensable in life sciences for visualizing cellular structures and dynamics in live specimens. Yet, conventional fluorescence imaging can suffer from phototoxicity, limited labeling efficiency, or perturbation of biological function. Label-free techniques such as interferometric scattering microscopy (iSCAT) offer a powerful alternative by detecting nanoscale structures based on their light scattering, without the need for dyes or genetic tags. iSCAT has enabled high-sensitivity detection of single proteins and viruses on clean surfaces. More recently, its application to live cells has been extended by using confocal illumination and detection, allowing suppression of out-of-focus light, yielding subcellular structures with high contrast. This development laid the foundation for biologically relevant label-free imaging. Here, we introduce interferometric image scanning microscopy (iISM). This next-generation technique combines interferometric detection with image scanning microscopy to achieve about 120 nm lateral resolution while operating at tenfold lower incident illumination power per diffraction limited spot, significantly reducing photodamage while enhancing signal-to-noise and contrast. Using iISM, we are able to visualize intracellular organelles such as the endoplasmic reticulum, actin cytoskeleton, mitochondria, and vesicles in live cells at essentially unlimited observation times. Importantly, iISM can be readily combined with confocal fluorescence microscopy, enabling correlation of label-free dynamics and structural information with molecular specificity. Our approach opens new avenues for studying dynamic biological processes, such as host-pathogen interactions, intracellular trafficking, or cytoskeletal rearrangements, under label-free, near-native conditions. iISM thus offers a powerful new tool for high-resolution, low-impact imaging of live cells, paving the way for new biological insights.

## Introduction

Light microscopy is indispensable in both material and life sciences and continues to advance toward higher spatial and temporal resolution, improved sensitivity, and enhanced imaging depth. In life sciences, live-cell imaging adds further demands of maintaining cell viability while minimizing phototoxicity and other perturbation effects.

Over the past two decades, fluorescence microscopy has entered the nanoscopic domain through super-resolution techniques, which surpass Abbe’s diffraction limit either based on stochastic single-molecule ((d)STORM[1, 2, 3], PALM[4]) or deterministic approaches (STED[5], SIM[6]). These methods leverage the molecular specificity of fluorescent probes to visualize sub-cellular structures with spatial resolution far below the diffraction limit, down to tens of nanometer (nm).

Even with diffraction-limited resolution, confocal laser scanning (fluorescence) microscopy (CLSM) remains a workhorse in cell biology due to its optical sectioning capability, which is an important property in particular for imaging three dimensional samples. In principle, CLSM can also enhance the lateral resolution beyond the conventional diffraction limit by a factor of 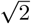 if the detection pinhole is closed below 0.2 Airy units (AU), and therefore is also often referred to as a super-resolution technique [7, 8]. In practice, however, this gain is rarely realized because most of the photons are discarded, leading to a substantial loss in signal-to-noise ratio (SNR). Image scanning microscopy (ISM), first formalized by Sheppard in the 1980s and experimentally demonstrated by Enderlein and coworkers in 2010 [7, 9], elegantly overcomes this trade-off. By replacing the single-element detector with an array detector, ISM recovers both the resolution of a closed pinhole and the SNR of an open pinhole, provided that the off-axis signals are reassigned to their correct positions (pixel reassignment) [10]. Variants of ISM have since been realized in both computational [10, 11, 12] and all-optical forms [13, 14, 15].

While fluorescence provides excellent molecular specificity, label-free methods are highly desirable for studying live cells to minimize or avoid perturbations. Interferometric scattering microscopy (iSCAT) [16, 17] has emerged as a powerful label-free technique capable of detecting nanoparticles, such as viruses [18], extracellular vesicles [19, 20] and single proteins [21, 22] with shot-noise limited sensitivity in the scenario of negligible background or clean surfaces. By interfering scattered light from a nanoscale object with a reference reflection at the sample–substrate interface, iSCAT achieves both exceptional sensitivity and broad applicability [23, 24]. However, imaging in live cells poses challenges due to the speckle created by coherent superposition of scattering events in complex heterogeneous media. To address this issue, recent work has extended iSCAT to imaging of intracellular structures in live cells using a confocal illumination and detection scheme, combining label-free contrast with optical sectioning in three-dimensional samples [25].

In this work, we propose and demonstrate the first experimental implementation of *interferometric* Image Scanning Microscopy (iISM). By combining the concepts of ISM and iSCAT, iISM enables high resolution of about 120 nm laterally, and label-free imaging inside live cells with substantially reduced illumination power. In order to adapt the ISM concept to an interferometric point-spread function (iPSF), we developed an adaptive pixel-reassignment (APR) algorithm tailored to coherent detection. Our new workflow restores high-resolution reconstructions with enhanced contrast-to-noise ratio (CNR) at about 10 times lower incident illumination power per diffraction limited spot compared to conventional confocal microscopy. We quantify with test objects, and then showcase the capabilities of iISM in live-cell imaging of intracellular organelles, including the dynamics of structures such as the endoplasmic reticulum. Furthermore, we demonstrate the complementarity of iISM and fluorescence ISM by correlative imaging of the actin cytoskeleton in fixed cells.

Our results establish iISM as a broadly applicable approach for minimally invasive, high-resolution imaging of living systems, combining the advantages of label-free interferometric contrast with the resolution and SNR gains from the ISM concept.

## Results

### Principle of interferometric ISM (iISM)

In this work, we developed a new ISM microscope that includes both interferometric scattering and fluorescence detection. The system employs a three-galvanometric mirror scanning module (Flimbee, Picoquant), controlled via the corresponding software (Symphotime 64, Picoquant), and a sCMOS camera (Orca-Fusion BT C15440, Hamamatsu) for detection (Fig. 1(a)). Camera acquisition is managed through open-source python-based software package *cam control* [26], that was modified to enable realtime iISM image acquisition and display (see Methods). Illumination is provided by a 445 nm diode laser (Cobolt MLD-06-01 150mW, Huebner Photonics) and spatially filtered through a polarization-maintaining single-mode fiber. The beam is separated from the detection path by a polarizing beam splitter and converted to circular polarization using a quarter-wave plate positioned before the objective. Circular polarization minimizes polarization-dependent scattering artifacts and improves interferometric detection efficiency. To reduce laser coherence artifacts and suppress intensity fluctuations, the laser driver current was continuously modulated during acquisition (see Methods, SI Fig. 7, 8). Additional experimental details are also provided in the Methods section. Both the light reflected at the coverglass interface and the light scattered by the sample are collected in reflection geometry through the same high-NA oil immersion objective, descanned by the galvanometric mirrors, and imaged in full-field on the sCMOS camera. The optical magnification was chosen to ensure significant oversampling of the interferometric point spread function (iPSF) (see below, and Methods). The detected intensity *I*_det_ corresponds to a confocal iSCAT measurement in reflection mode and can be expressed as:

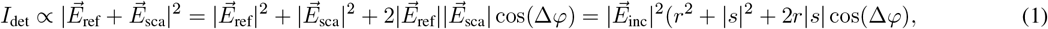

where 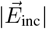 is the incident electric field amplitude, *r* is the reflectivity at the coverglass imaging medium interface, and |*s*| is the scattering amplitude of the object [16, 17]. In a confocal geometry, the interference occurs between two quasi-spherical waves and the relative phase between reflected and scattered electric fields is:

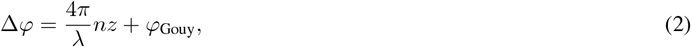

with *n* the refractive index of the medium, *z* the axial position of the scatterer relative to the interface, *λ* the illumination wavelength, and *φ*_Gouy_ the Gouy phase [17, 25, 27].

**Figure 1.**
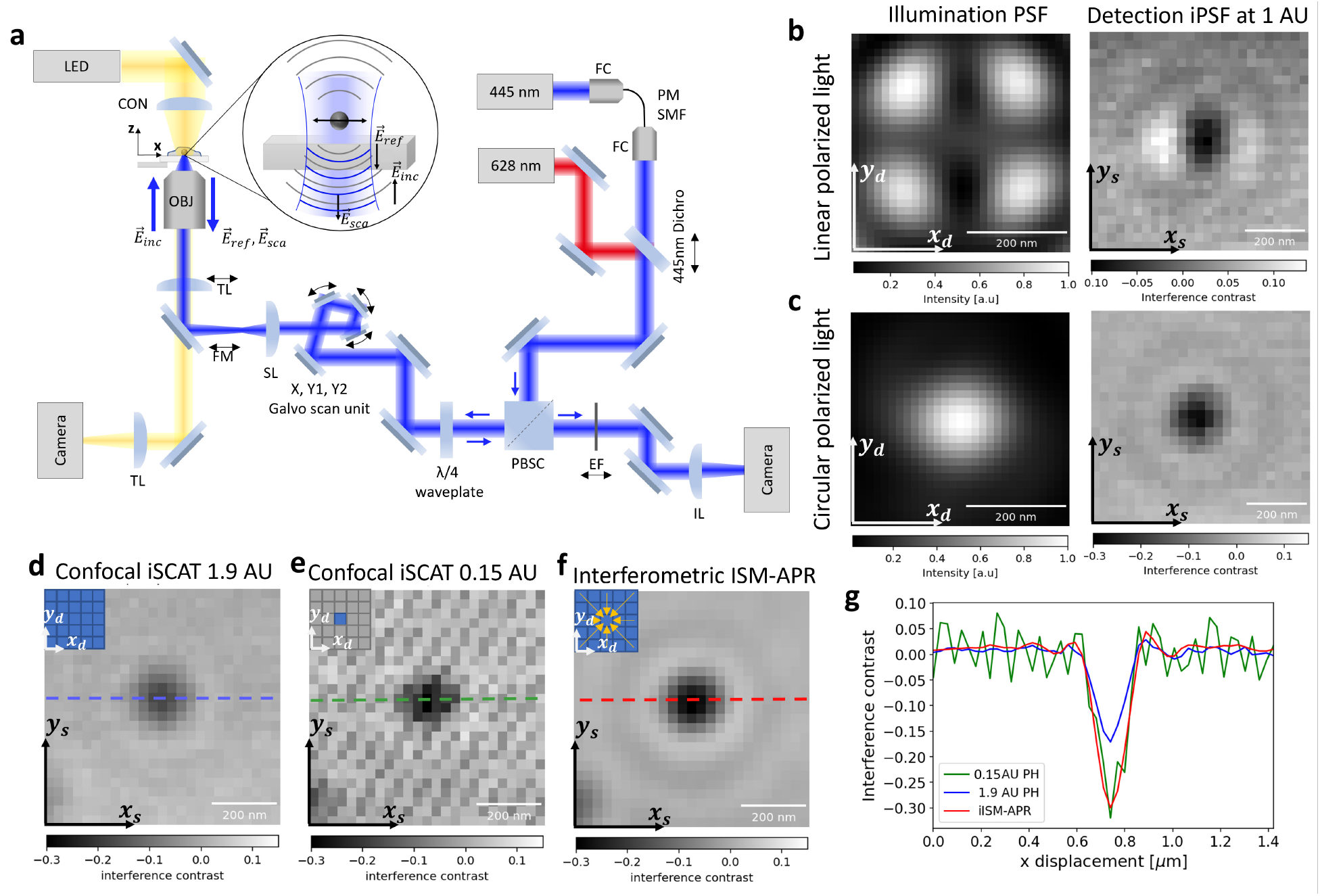
Interferometric ISM (iISM) principle and interferometric point-spread-function (iPSF). **(a)** Schematic of the image scanning microscope. OBJ objective, TL tube lens, FM flip mirror, SL scan lens, PBSC polarizing beam splitter cube, EF emission filter, IL imaging lens, CON condensor, FC fiber coupler, PM SMF polarization-maintaining single-mode fiber. Black arrows indicate mechanical motion degrees of freedom. Side-by-side comparison of the illumination PSF and detection iPSF at 1 AU of a 60 nm polysterene nanoparticle for **(b)** linear polarized light and **(c)** circular polarized light. The coordinates *x*_*d*_, *y*_*d*_ denote the camera sensor plane and *x*_*s*_, *y*_*s*_ the sample plane, respectively. **(d)** Resulting iPSF from a corresponding open pinhole and **(e)** closed pinhole confocal iSCAT image (for details see main text). **(f)** Resulting iPSF from iISM after adaptive pixel-reassignment (APR), with same incident illumination power and number of detected photons. **(g)** Line profiles of the iPSF in the three configurations as indicated in (d-f) at 1.9 AU (blue), 0.15 AU (green) and after iISM-APR (red). Airy unit (AU). All scale bars are 200 nm.

In conventional ISM, the incoherent nature of fluorescence imaging renders the effective PSF as the product of illumination and detection intensity PSFs, i.e. *I* = *obj* ⊛ (*PSF*_*det*_ · *PSF*_*ill*_), where *obj* denotes the object function and ⊛ is the convolution operator [28, 29]. However, under coherent conditions, as in iISM, the detected intensity of an object is a function of the amplitude PSF *h*:

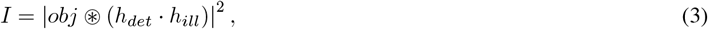

with *h*_*det*_ and *h*_*ill*_ denoting the detection and illumination amplitude PSFs, respectively [28, 29, 30]. A full theoretical description of the iPSF is beyond the scope of the present work, however for the following it suffices to note that the phase is directly encoded in the detected intensity iPSF, which we are going to account for by introducing a modified adaptive pixel-reassignment (APR) workflow as described in the following section.

To acquire an iISM dataset, at each position of the galvo scanner in the sample plane 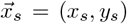, a microimage 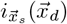 is recorded with the detector pixel positions 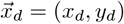. This procedure results in a four-dimensional dataset. Typically, the size of a microimage corresponds to about 1.4 Airy units (AU, with 1 AU = 1.22*λ/*2NA), which for our parameters (*λ* = 445 nm, NA = 1.4) corresponds to 9 effective camera pixels (see Methods for details). To recreate an open-pinhole confocal iSCAT image [25], the detected intensity at each sample position is obtained by summing all intensity values of the corresponding microimage, 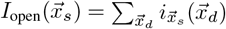. In contrast, for a closed-pinhole confocal iSCAT image we evaluate only the central pixel of each microimage, in direct analogy to the fluorescence ISM concept (see Methods).

The illumination and detection PSFs together determine the imaging properties of the confocal microscope (see eq. (3)), and are therefore critical for optimizing resolution and contrast. Unlike fluorescence ISM or confocal iSCAT, iISM provides simultaneous access to both PSFs, which we exploit to control the polarization and optimize the effective iPSF shape. For calibration, we used sparsely distributed 60 nm polystyrene nanoparticles immobilized on a cover glass and mounted in PBS (see Methods), and acquired a z-stack with 29 nm voxel size (see SI Figs. 1, 2, and 4).

Figures 1(b,c) compare the effects of linear and circular polarization on the illumination PSF and the resulting iPSF, demonstrated by evaluating a single 60 nm nanoparticle. The illumination PSF was estimated by recording the reflection as a function of the beam position of the mostly empty coverglass and averaging the intensity of the microimages across all scan positions, 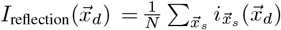, with *N* the total number of scan positions. The corresponding detection iPSF was obtained by reconstructing an open-pinhole confocal iSCAT image of a single nanoparticle and applying background normalization to determine the interference contrast (see Methods). For linear polarization (Fig. 1(b), left), by looking closely, one sees that the illumination PSF is elongated along the polarization axis, consistent with theoretical predictions for tightly focused Gaussian beams at dielectric interfaces under high-NA conditions [31]. The corresponding detection iPSF (Fig. 1(b), right) also shows a clear elongation along the sample’s x-axis. Moreover, iISM datasets acquired with linearly polarized light reveal several phase flips across different off-axis pinhole positions, caused by the non-uniform phase distribution of the illumination PSF (see SI Fig. 3). Ideally, however, an iISM dataset of off-axis iPSFs should exhibit constant phase and rotational symmetry to enable robust pixel reassignment.

To address this, we introduced a polarizing beam splitter cube and a quarter-wave plate to generate circularly polarized light in the sample plane. The quarter-wave plate was adjusted to maximize PSF isotropy under stationary beam conditions, accounting for potential anisotropies introduced by the optical components in the beam path. With circular polarization, the illumination PSF (Fig. 1(c), left) clearly exhibits improved rotational symmetry. The corresponding detection iPSF (Fig. 1(c), right) is substantially more isotropic, independent of virtual pinhole size or off-axis pinhole position (see SI Fig. 3). This ability to directly measure and optimize the illumination PSF without additional wavefront sensors highlights a key advantage of iISM.

Figures 1(d,e) compare the iPSFs under circular polarization for open-pinhole and closed-pinhole confocal iSCAT configurations after background normalization. As expected, the closed pinhole iPSF exhibits a full width at half maximum (FWHM) of about 122 nm *±* 5 nm (see SI Fig. 1), in good agreement with the theoretical value for a closed pinhole for our parameters FWHM_theo_ ≈ 0.4*λ/*NA ≈ 127 nm [8]. However, this comes at the cost of an increased noise floor due to reduced number of detected photons. To quantify both interference contrast and noise floor, we calculate the contrast-to-noise ratio (CNR), where we define the Noise Equivalent Contrast (NEC) as the standard deviation of pixel intensities within a sample-free region of interest (ROI) (see Methods and SI Fig. 2). For the closed-pinhole configuration, we measure a maximum negative interference contrast of about 0.3 (Fig. 1(g), green) at NEC_closed_ = 0.03, yielding CNR_closed_ = 10. In the open-pinhole case, the maximum negative interference contrast is reduced to 0.15 (about 50% lower), but the noise floor is also reduced to NEC_open_ ≈ 0.011, resulting in a slightly improved CNR_open_ ≈ 14.

Using iISM with our modified APR algorithm (detailed in the next section), we obtain a lateral resolution limit of FWHM = 120 nm *±* 4 nm (see SI Fig. 1) and the doubled contrast of the closed pinhole configuration (Fig. 1(g), red), while achieving the lowest noise floor of NEC_iISM_ ≈ 0.008. This yields a significantly improved CNR_iISM_ of about 38, almost four times higher than the closed-pinhole case and about three times higher than the open-pinhole case. This superior CNR, combined with the enhanced resolution, establishes iISM as the most sensitive and highest-resolution method among the three configurations, and is analogous to the “super-concentration of light” effect reported in fluorescence ISM [32].

### iISM with adaptive pixel-reassignment (APR)

Fluorescence ISM achieves super-resolved reconstructions by combining signals from an array of off-axis detector elements and computationally or optically reassigning them to their correct spatial positions, thus utilizing the additional information to improve the PSF. The most widely used approach is pixel-reassignment (PR), where the shift vectors 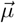 are estimated from the detector geometry. A simple rule is to apply a constant PR factor of e.g. 1*/*2 of the pixel displacement, which assumes the same wavelength for both illumination and detection light [7]. Historically, this approximation was also applied to fluorescence ISM, even though the underlying emission process is incoherent and therefore does not strictly justify the same reassignment factor due to Stokes-shifted fluorescence emission [9, 10]. To address this mismatch, adaptive pixel-reassignment (APR) was introduced, which estimates optimal shift vectors directly derived from the measured PSFs [11]. In fluorescence ISM, APR has been shown to yield improved resolution and SNR, while it intrinsically also accounts for experimental imperfections, detector misalignments, and deviations from the ideal 1*/*2 rule. The situation is more subtle for iISM, since the detected signal is coherent and therefore encodes both amplitude and phase. In this case, one has to explicitly consider the phase distribution of the iPSF before reassignment can be applied. For circularly polarized, tightly focused Gaussian beams, the illumination phase in the focal plane is known to be nearly flat (wavefront curvature negligible) up to approximately ~1 AU lateral radius [33, 31]. This motivates our choice of using a radius of ~1 AU as an upper limit for the region in which our new iISM-APR algorithm is applied. To adapt APR to the interferometric case, we modified the standard workflow to take into account the phase information of the iPSF as follows. Specifically, we use the radial variance transform (RVT) [34], which converts an interferogram into an intensity-only map that reflects the local degree of symmetry. This procedure effectively emulates the incoherent fluorescence PSFs used in conventional APR, while still preserving the essential spatial information from the interferometric detection. In effect, the phase-correlation of the individual RVT images with the central pixel image defines the required shift (see Fig. 2 (e)). The resulting RVT-transformed maps of both on-axis and off-axis pinholes are then supplied as inputs to the APR algorithm (see Fig. 2 (b)). For the implementation of APR, we took advantage of the readily available *Brighteyes* APR python library [35], which allowed us to perform automated image registration on the obtained RVT dataset. From these registrations, we obtained a set of shift vectors that originate from our interferometric data (Fig. 2 (e)). Finally, these RVT-APR shift vectors were applied back to the original iISM dataset, yielding reconstructions with enhanced spatial resolution. Importantly, this workflow enabled us to achieve a superior CNR at about 10 times lower incident illumination power, which is particularly advantageous for live-cell imaging as we will describe in the next section.

**Figure 2.**
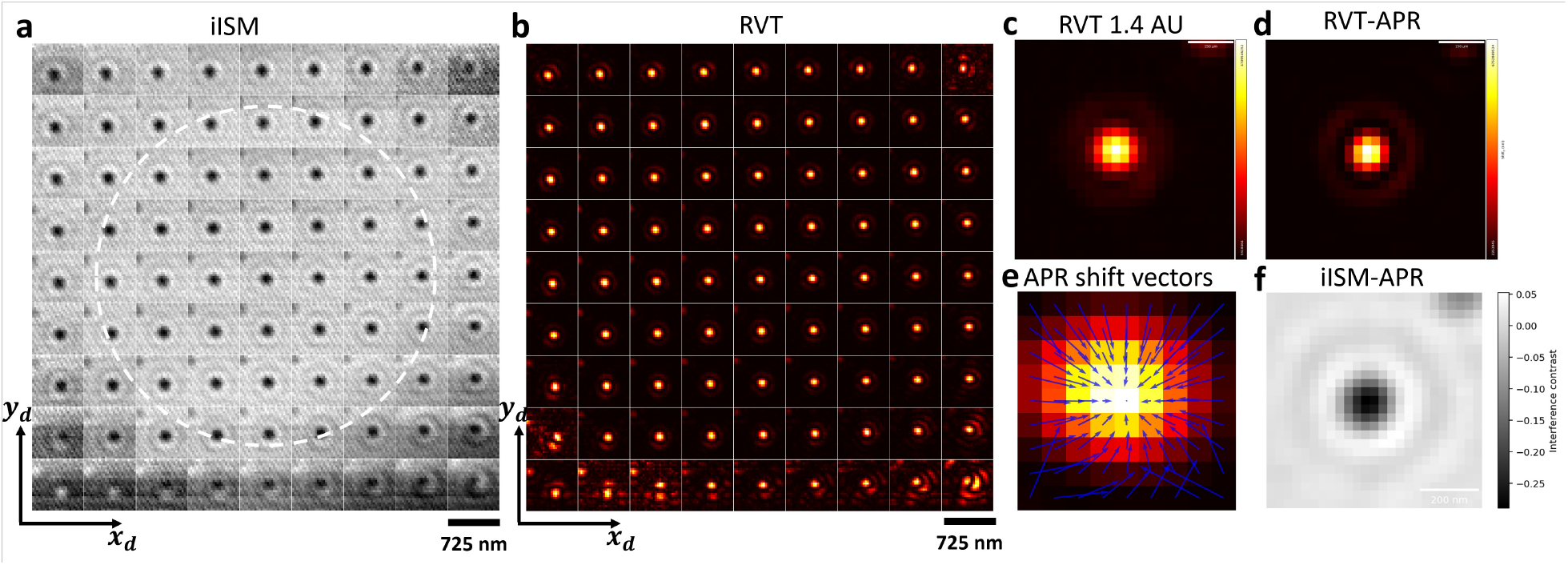
Interferometric ISM (iISM) with adaptive pixel-reassignment (APR) workflow. **(a)** Matrix representing the scanned interference images of a single 60 nm polystyrene nanoparticle (PNP) with a 9×9 array detection. Each position corresponds to a closed off-axis virtual pinhole of 0.15 AU with respect to the central pinhole. White dashed circle corresponds to 1 AU. The full area of 9×9 pixels corresponds to 1.4 AU. **(b)** Matrix, representing the radial variance transform (RVT) of (a). Scale bar (a,b) 725 nm. **(c)** RVT image of a single 60 nm PNP with open pinhole (1.4 AU) derived as the sum from matrix (b). **(d)** RVT image corresponding to (c) after applying APR. **(e)** Fingerprint map with the superimposed shift vectors (blue) from APR based on the RVT matrix in (b). **(f)** Resulting iISM image of a single iPSF after applying the obtained shift vectors from (e) to the iISM matrix in (a). Scale bar 200 nm.

### iISM imaging of intracellular organelles in live cells

To assess the performance of iISM under physiologically relevant conditions, we applied the method to live-cell imaging of COS-7 cells, focusing on intracellular organelles such as mitochondria, the endoplasmic reticulum (ER), vesicles, and the actin cytoskeleton. Figure 3 (a) shows an overview of a single optical section with a size of about 40 times 80 *µ*m, acquired at an incident illumination power of about 0.5 *µ*W, enabling virtually unlimited observation time without inducing visible photodamage or compromising cellular integrity. Major organelles, including the nucleus (N), mitochondria (M), ER, vesicles (V), actin cytoskeleton (A), plasma membrane, and lamellipodia (L), are readily distinguishable, despite the absence of labels. Notably, individual structures exhibit positive and negative interference contrast (see Figs. 3 (b,c)), indicative of small axial displacements relative to the optical section, highlighting iISM’s intrinsic sensitivity to nanoscale three-dimensional morphology and displacements (see eq.(2)).

**Figure 3.**
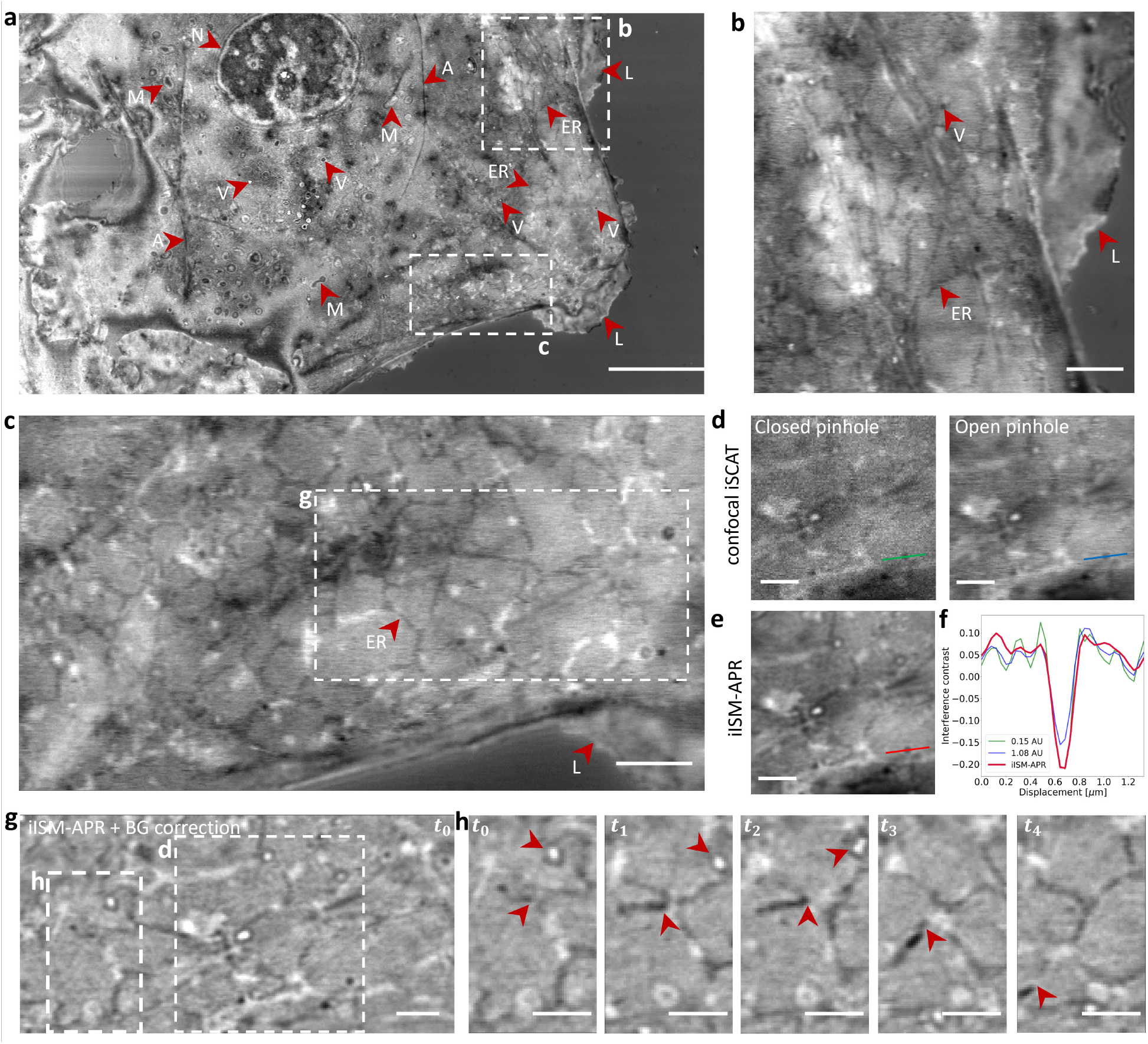
Live-cell interferometric ISM captures dynamics of intracellular organelles with enhanced contrast and resolution at about 0.5 *µ*W incident illumination power. **(a)** Overview of a COS-7 cell. Scale bar 10 *µ*m. Arrows mark individual organelles that are visible in this particular optical section. Nucleus (N), mitochondria (M), actin cytoskeleton (A), plasma membrane and lamellipodum (L), the endoplasmic reticulum (ER) as well as vesicles (V). Notably, positive and negative contrast of the same type of organelle indicate slightly different axial positions with respect to the optical section. **(b)** Upper close-up from (a) showing ER, vesicles and lamellipodium. Scale bar 2 *µ*m. **(c)** Lower close-up from (a) showing ER and lamellipodium. Scale bar 2 *µ*m. **(d)** Confocal iSCAT evaluated for the close-up of (g) as indicated with white dashed square. (Left) Closed pinhole and (right) open pinhole analysis. **(e)** iISM-APR of the same region as (d). **(f)** Line profiles of interference contrast of vesicle cross-section as indicated in (d,e) for closed pinhole (green), open pinhole (blue) and iISM-APR (red). **(g)** iISM-APR timelapse of close-up in (c) after additional background correction to increase visibility of organelles on flat-fielded background (see Methods). **(h)** Timelapse of close-up in (g) with exemplary four consecutive time points (*t*_0_ − *t*_4_) at time increments of 10s. Upper red arrows indicate vesicle motion, lower red arrows indicate ER remodelling. Scale bars (d-h) 1 *µ*m.

To benchmark the performance against conventional confocal iSCAT, we compared in Figure 3 (d) the same square region of interest from panel (g) reconstructed with a closed pinhole and an open pinhole (as described in Methods). While the closed pinhole provides improved lateral resolution, it suffers from increased noise, which typically restricts its use in dynamic imaging at low incident laser powers. Conversely, the open pinhole improves photon collection but sacrifices both resolution and contrast, leading to blurred organelle boundaries. Our iISM approach overcomes these limitations by combining confocal interferometric detection with APR, thereby recovering high spatial resolution while increasing CNR (Fig. 3 (c)). Quantitative line profiles across a vesicle (Fig. 3 (f)) demonstrate a clear enhancement in both CNR and lateral resolution for iISM-APR (red) compared to the other two configurations.

For further analysis of dynamics, a flat-field background correction (as shown in [25], and Methods) was applied to the APR-reconstructed image, enabling visualization of organelles with increased clarity against the residual interferometric background (Fig. 3 (g)). This approach allowed us to track vesicle motion and ER remodeling over extended periods at seconds temporal resolution (10 s frame interval shown here). Note that the achieved temporal resolution here is limited by the maximum framerate of the camera, and does not have a photophysical upper limit like in fluorescence ISM. The trajectories of individual vesicles (upper arrows) and dynamic reorganization of ER tubules (lower arrows) underscore the ability of iISM to resolve and follow intracellular dynamics in real time.

### Correlative iISM and fluorescence ISM of the actin cytoskeleton

To further validate the structural information obtained with iISM and to assess its complementarity to fluorescence imaging, we performed correlative experiments on fixed COS-7 cells, labeling the actin cytoskeleton with Phalloidin-Alexa Fluor 647 (AF647). For fluorescence excitation, we combined the output of a 628 nm fiber laser (F-04306-106, 1W, MBPC) with an edge shortpass filter (457nm, FF457-SDi01-25×36, Semrock) with the beam of the 445 nm laser, and ensured that the reflections at an empty coverglass region of both lasers are concentric on the camera. In order to detect fluorescence emission we then sequentially imaged with 445 nm for iISM and 628 nm for fluorescence ISM, where we inserted a long-pass emission filter (633 nm, razor edge, LP02-633RE-25, Semrock) in the detection path (see Fig. 1 (a)) to spectrally separate the fluorescence emission from the excitation.

Figure 4 (a) shows the label-free iISM image of an actin-rich lamellipodium region after background normalization, where extended filamentous structures are clearly visible without the need for staining. The corresponding fluorescence ISM reconstruction of phalloidin-AF647 (Fig. 4(b)) provides specific labeling of the actin network. Overlay of the two modalities (Fig. 4(c)) confirms the excellent spatial correspondence between actin filaments detected by iISM and those revealed by fluorescence, demonstrating that interferometric scattering directly reports on filamentous cytoskeletal structures. A magnified view of the boxed region (Fig. 4(d–f)) highlights the level of detail provided by iISM in comparison to fluorescence ISM. Both modalities capture the same filamentous bundles (cyan arrowheads), while iISM additionally reveals nanoscale contrast variations along filaments and adjacent unlabeled structures that remain invisible in fluorescence. Notably, due to the scattering background from other structures in between the actin bundles, the finer actin mesh is mainly visible in fluorescence ISM. This underlines the complementarity of the two techniques: fluorescence provides molecular specificity, whereas iISM extends the accessible information by reporting on unlabeled structures, such as vesicles and focal adhesions. To quantify the correspondence between modalities, we extracted line profiles across individual filaments (Fig. 4(g)). The fluorescence channel (red) identifies the actin filament position, while the iISM channel (black) resolves fine contrast modulations arising from nanoscale variations in scattering cross-section and axial position. The profiles illustrate that iISM achieves co-localization with fluorescence at sub-diffraction accuracy, while providing additional axial sensitivity through phase variations. Together, these results establish iISM as a powerful label-free complement to fluorescence ISM, enabling correlative imaging of cytoskeletal architecture and beyond.

**Figure 4.**
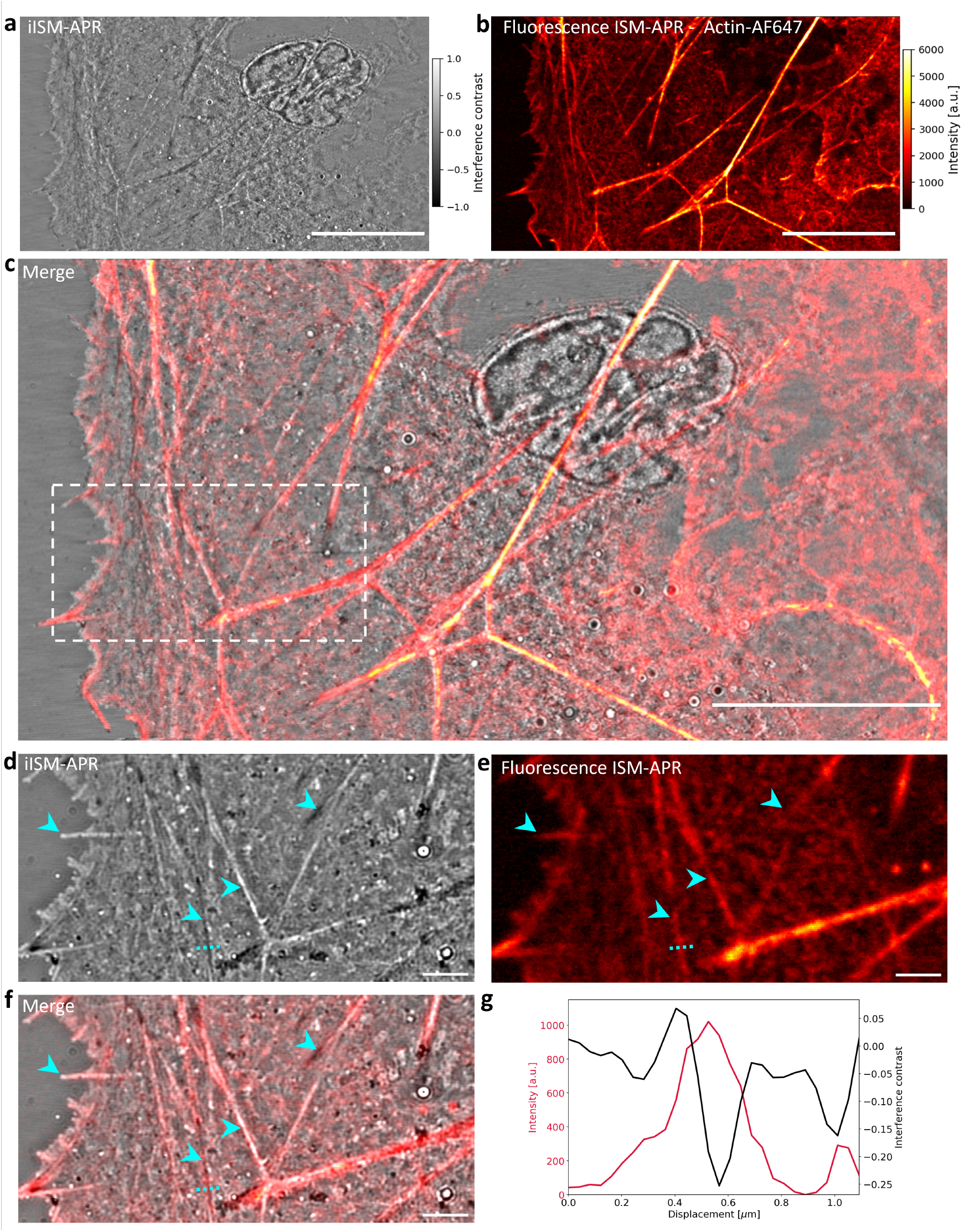
Correlative iISM and fluorescence ISM of actin cytoskeleton in fixed COS-7 cells. **(a)** Label-free iISM image of the cell, revealing extended filamentous actin structures. **(b)** Fluorescence ISM reconstruction of Alexa Fluor 647-labeled actin of the same field of view. **(c)** Merge of (a) and (b) demonstrates spatial correspondence between modalities. Scale bars (a-c) 15 *µ*m. **(d)-(f)** Close-up of white dashed box in (c) in both modalities. Cyan arrows (from left to right) highlight the tip of a filopodia protruding from the cell, two actin fiber bundles with opposing contrast due to their relative axial position, and a focal adhesion. Scale bars (d-f) 2 *µ*m. **(g)** Line profiles across cyan dashed line in (d-f) showing co-localization of actin fluorescence (red) and iISM (black) interference contrast.

## Discussion and Outlook

The iISM approach introduced here demonstrates that coherent scattering signals can be harnessed within the ISM framework to achieve both high resolution and high sensitivity. By combining interferometric detection with a modified APR algorithm based on RVT, we realized a lateral resolution of about 120 nm, quantified by the FWHM of the iPSF, while obtaining a noise reduction of up to an order of magnitude compared to conventional detection. This CNR enhancement translates directly into higher contrast for objects of identical scattering cross-sections, reminiscent of the “super-concentration of light” effect described in fluorescence ISM [32].

The implementation in live cells further highlights the potential of iISM as a label-free modality for imaging intracellular organelles. Importantly, here we have demonstrated maintaining the resolution of a closed pinhole confocal iSCAT with only about 0.5 *µ*W incident illumination power in the focused illumination spot, which is about 10 times lower than previously reported [25]. An improvement of one order of magnitude laser power reduction is significant for potential applications in primary cells, where the phototoxicity threshold is typically even lower than in immortalized model cell lines. Furthermore, the correlative experiments with fluorescence ISM establish iISM as a powerful complement rather than a replacement, enabling side-by-side evaluation of molecularly labeled and label-free structures. This opens new possibilities for hybrid strategies, where fluorescence provides molecular specificity while iISM delivers structural contrast, context, and quantitative scattering information.

Beyond our implementation, the method is highly adaptable. The camera can be trivially exchanged for a SPAD array, which would provide higher temporal resolution analogous to recent fluorescence ISM advances [11]. Parallelized detection schemes, as described for fluorescence ISM [13] or for confocal iSCAT [25, 36], could further accelerate imaging, extending iISM to dynamic processes at the millisecond scale. In addition, computational staining approaches, where segmentation and classification can substitute for chemical labeling, suggest that iISM may deliver molecular specificity through purely algorithmic means. This concept aligns with recent demonstrations of segmenting endoplasmic reticulum morphology from label-free scattering data by training of a neural network on fluorescence and scattering image pairs [25]. The recent publication of coherent anti-Stokes Raman ISM underlines the versatility of the principle and suggests that other coherent contrast mechanisms could be integrated into iISM [37].

Notably, iISM can be readily integrated into already existing commercial confocal fluorescence ISM systems, such as Airy scan from Zeiss [38], Nsparc from Nikon [39], and Luminosa from Picoquant [40], by substituting the main dichroic mirror with a polarizing beam splitter and adding a quarter-wave plate for circular polarized light at the desired wavelength. Looking ahead, a combination of iISM with single-molecule fluorescence ISM (SM-ISM) [41] promises new insights into cellular architecture by uniting ultrasensitive scattering detection with molecular specificity. We anticipate that such hybrid strategies will not only deepen our understanding of cell structure and function but also establish iISM as a cornerstone method within the broader ISM family.

## Methods and Materials

### Microscope setup

For this work, we built a custom ISM microscope that enables both interferometric scattering and fluorescence detection as shown in Figure 1 (a). The incident illumination beams are provided by a 445 nm diode laser (Cobolt MLD-06-01 150mW, Huebner Photonics) and 628 nm fiber laser (F-04306-106, 1W, MBPC). We control the power of the 445 nm laser by modulating its driver current with a sine wave of peak-to-peak amplitude *V*_pp_ = 50 mV, offset of *V*_off_ = 100 mV, and at 150 kHz modulation frequency. The incident illumination power was adjusted with a variable neutral density (ND) filter wheel, such that the power at the sample plane was 0.5 *µ*W. The 445 nm laser output is fiber coupled into a polarization maintaining single-mode fiber for spatial filtering. The 445 nm and 628 nm laser beams are combined via a shortpass filter (457nm, FF457-SDi01-25×36, Semrock). The illumination and detection path are separated by a polarizing beam splitter cube (PBSC) followed by a zero-order quarter wave plate (WPQ10M-445, Thorlabs) to yield circular polarized light. For spectrally separating the fluorescence emission from its excitation in the correlative iISM and fluorescence ISM measurement, an additional long-pass emission filter (633 nm, razor edge, LP02-633RE-25, Semrock) is flipped in front of the camera. The circularly polarized 445 nm light is partially reflected at the cover glass-imaging medium interface to provide the reference beam, and the partially transmitted light is scattered by the sample depending on the scattering cross-section of the e.g. nanoparticle or cellular structure. Both the scattered and reflected light are collected in a reflection geometry and homodyne detected on the camera, which gives rise to the detection intensity in eq. (1). The laser beams are steered via a three-galvanometric scanning mirror system (Flimbee, Picoquant). The Flimbee scanning system allows precise positioning of the incident light beam pivot point in the back-focal plane of the objective, improving field homogeneity in the sample plane during scanning. The Flimbee is controlled via the combination of the Flimbee control unit (Picoquant), and a stand-alone Time-Correlated Single Photon Counting (TCSPC) system (Picoharp 300, Picoquant), which is required for satisfying the hardware initialization routine of the Flimbee system and the corresponding control software (Symphotime 64, Picoquant). The Flimbee system is mounted on a joint baseplate with a commercial microscope stand (IX73, Olympus Evident Scientific), which is equipped with a high-NA oil immersion objective (Uplan SApo 100x 1.4 NA oil immersion, Olympus) mounted on a manual rotation nosepiece with z drive for coarse focusing. The sample is mounted in the sample holder of a piezo-driven precision stage (PI nano XYZ P-545 stage with E-727 digital servo controller, Physik Instrumente) enabling precise x,y positioning of the sample and z scanning. The microscope stand is further equipped with an LED light source and condensor, to allow for brightfield transmission imaging onto a second camera (U3-3270CP-M-GL, IDS) that we attached to the left port. We use this auxillary modality for orienting and positioning the sample before confocal scanning, in particular during cell experiments for finding specific areas of interest. In order to couple the confocal beam path into the microscope stand’s optical main axis, a custom adapter provided from Picoquant is equipped with a flip mirror and tube lens (focal length 180 mm). The scan lens (focal length 90 mm) is mounted in the Flimbee housing and together with the tube lens projects the pivot point of the back focal plane of the objective onto the scan mirror system. Both fluorescence and interferometric scattering are collected by the same objective in a reflection geometry, and then descanned by the same scan mirror system before the imaging lens (focal length 400 mm, AC254-400-A, Thorlabs) conjugates the sample plane onto the the main camera (Orca-Fusion BT C15440, Hamamatsu) with an overall magnification of 440x. With a physical pixel size of 6.5 *µ*m the effective optical pixel size on the camera sensor amounts to 15 nm in the sample. Considering the illumination wavelength of 445 nm and NA of 1.4, 1 AU = 194 nm, i.e. one pixel on the camera corresponds to about 0.08 AU in the sample plane. For further analysis of the datasets we then binned the camera pixels 2×2, which then yields an effective optical pixel size of 30 nm and one camera pixel corresponding to 0.15 AU in the sample plane.

### Hardware control and data acquisition

For synchronization of the hardware components, the pixel clock and line clock output signals from the Flimbee scan control unit were combined using a logic AND gate (DM74LS08, Fairchild semiconductor). The resulting signal was used as an external hardware trigger for the camera, ensuring that exactly one frame was acquired per scan position in the sample plane. Data acquisition was carried out using a custom *cam-control* software based on *pyLabLib* (developed by Alexey Shkarin, https://github.com/SandoghdarLab/pyLabLib-cam-control [26]). To enable ISM measurements, we modified the open-access cam-control software by implementing an ISM plugin (that can be downloaded from our repository). The modified software allows live display of ISM images with an effectively open pinhole during acquisition. The galvanometric scanning mirrors were controlled via Symphotime 64 (Picoquant) and served as master for the data acquisition.

For each iISM dataset, we acquired different sized ROIs on the camera chip, depending on the specific measurement at hand, since the ROI size on the camera determines the maximum read-out speed and with this, the effective frame rate of imaging the sample. Specifically, we used the following acquisition parameters for the datasets shown in the corresponding figures: Fig. 1,2: 29 nm scanning pixel size (px size), 256×128 px, 300 *µ*s dwell time, camera exposure time 0.15 ms. Fig. 3: a) b) 39 nm px size, 2048×1024 px, 250 *µ*s dwell time, camera exposure time 0.11 ms. c) 39 nm px size, 512×256 px, 140 *µ*s dwell time, about 37s frame period, camera exposure time 0.07 ms. d) e) g) h) 39 nm px size, 256×128 px, 140 *µ*s dwell time, about 10s frame period, camera exposure 0.07 ms. Fig. 4: a) 78 nm px size, 1024×512 px, 250 *µ*s dwell time, camera exposure time 0.11 ms, b) 78 nm px size, 1024×512 px, 500 *µ*s dwell camera exposure time 0.380 ms exp, d) 39 nm px size, 512×256 px, 250 *µ*s dwell time, camera exposure time 0.11 ms.

### Data analysis

#### Confocal measurement

In order to obtain the corresponding confocal measurement from an interferometric or fluorescence ISM dataset, we either evaluate the single central pixel in the detector plane 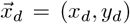 for each scan position in the sample plane 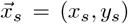 corresponding to an effectively closed pinhole confocal, or sum over all detector pixels, corresponding to an open pinhole confocal geometry:

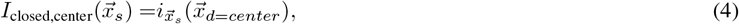

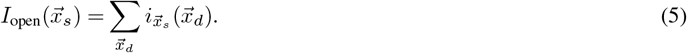

The results are equivalent to either a confocal iSCAT or confocal fluorescence measurement, respectively, with open or closed pinhole.

#### Contrast

Optical contrast describes the ability to distinguish the optical response of an object from the background *I*_bg_ detected when the object is not present. In iSCAT, the reference electric field typically serves as the background of the detected intensity, such that the interference contrast can be defined as:

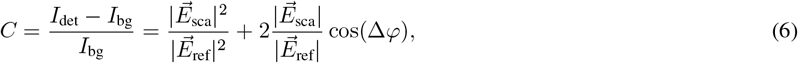

with the reflected electric field 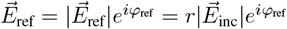 and the scattered electric field 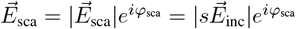. The maximum positive and negative interference contrast values are obtained for Δ*φ* ∈ *{*0, *π}*, yielding:

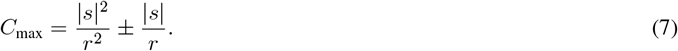

For practical considerations, we estimate the background intensity *I*_bg_ by low-pass filtering each iISM frame using a Gaussian filter with a typical sigma of 9 pixels in the sample plane (unless stated otherwise), corresponding to a FWHM of about 830 nm depending on the effective optical pixel size. This procedure effectively flat-fields the detected intensity, allowing contrast measurements of intracellular structures relative to their local environment rather than to the glass substrate. In case of the iPSF measurements of sparsely distributed nanoparticles we extracted the interference contrast by calculating the background intensity as the median of the whole image evaluated either in iISM-APR, closed-pinhole, or open-pinhole configuration, respectively.

#### Contrast-to-noise ratio (CNR)

As a metric to quantify the image quality, we consider the contrast-to-noise ratio (CNR) as:

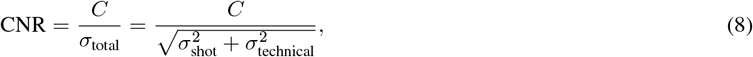

where *σ*_total_ is the total noise consisting of *σ*_shot_, which represents the inherent photon shot noise, and *σ*_technical_, which accounts for technical noise sources, such as scanning mirror noise, sCMOS camera read noise and fixed-pattern noise, as well as technical laser intensity fluctuations above shot noise.

#### Noise Equivalent Contrast (NEC)

To quantify the total noise floor in an image, we computed the Noise Equivalent Contrast (NEC) in a sample-free region of interest (ROI):

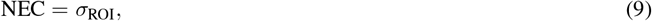

where *σ*_ROI_ is the standard deviation of the pixel intensities. This metric provides a single-value estimate of the effective noise floor in each imaging configuration (open/closed pinhole, iISM-APR), independent of sample structure. Lower NEC values indicate reduced noise variance. Unlike the CNR, which requires a defined signal, NEC strictly characterizes the noise contribution of the imaging system in the three different configurations.

#### iISM analysis

For each iISM dataset, we acquired microimages with an effective area on the camera of at least 1 AU. Each pixel on the camera thereby corresponds to an individual closed “virtual” pinhole of 0.15 AU either on or off the optical axis of the ISM detector array. From these microimages, we generated an iISM pinhole stack of single-pixel pinhole images according to eq.(4), which serves as the basis for subsequent analysis.

#### Radial variance transform (RVT)

To enable robust registration of the iISM pinhole stack independent of the interferometric phase, we applied RVT to it. RVT computes, for each pixel, the variance of intensity values along concentric circular areas of increasing radius and generates a new image in which pixel intensity encodes the degree of radial symmetry (see Fig. 2 (b) and SI Fig. 1 (b)). This approach exploits the fact that centers of radial symmetry can be identified by a low mean of variance combined with a high variance of means of pixel values across different radii, enabling robust identification of symmetry centers even in noisy data or the presence of asymmetric iPSFs [34]. Importantly, the RVT output no longer carries phase modulations from interferometric contrast and can therefore be used for subsequent pixel-reassignment analysis. For our data, we used RVT radii of *r*_min_ = 1 px, and *r*_max_ = 4 px to obtain the corresponding RVT pinhole stack for further analysis.

#### Adaptive pixel-reassignment (APR)

APR was performed using image registration based on phase correlation of the off-axis raw images with respect to the central one, as detailed in [35]. Image registration of the RVT pinhole stack yielded shift vectors 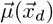 for each off-axis pinhole image relative to the central one. These vectors represent the spatial displacement of the corresponding effective detection iPSF relative to the optical axis. The obtained shift vectors were then applied to the original iISM pinhole stack, enabling precise alignment of the off-axis pinhole images prior to summation. The final iISM-APR image was calculated as the sum of the aligned iISM pinhole stack:

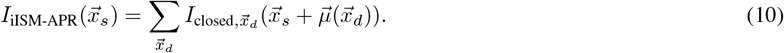

For three-dimensional (3D) datasets, either *xy* and *z* (e.g. iPSF SI Fig. 4) or *xy* and time (e.g. live-cell timelapse Fig. 3), the iISM-APR algorithm was applied to a single reference plane (central focal plane) or reference time point (first frame), and the resulting shift vectors were subsequently applied to the entire 3D dataset. For fluorescence ISM in Figure 4 (b), the conventional APR algorithm was applied as described in [35].

#### Software packages

All data analysis was performed using custom code written in Python 3.11.7 using standard Python libraries as well as scipy.ndimage v1.11.4, imgrvt v1.0.0 [34], trackpy v0.6.3 [42], and brighteyes-ism v1.3.4 [35].

### iPSF calibration measurements

For calibration measurements of the iPSF, 60 nm polystyrene nanoparticles (Nanospheres NIST, size 59 *nm ±*3 *nm*, Cat. No. 3060A, Lot. No. 289283, Thermo Fisher) were diluted 1:100 in mQ. Cover glasses (high precision, 22×22 mm No. 1.5H 170 *µm ±* 5 *µm* thickness, VWR) were cleaned by sequential rinsing with ethanol and deionized water before drying with N_2_. Circular silicone spacers (Cat. No. 70326-56, size 2.5mm, volume 150 *µ*l, EMS) were placed on top of the cleaned cover glass to form a chambered sample mount. In order to immobilize the nanoparticles on the surface of the cover glass, the cover glass was incubated for 5 minutes with poly-L-lysine (P8920, Simga Aldrich) to impart a net positive surface charge, after that the excess solution was removed. The nanoparticle solution was added to the cover glass chamber and after incubation for 5 minutes, the excess solution was removed and replaced by PBS, which then served as the mounting medium for imaging. The sample chamber was sealed with an additional cover glass (18 mm round, VWR) to avoid evaporation, which was cleaned beforehand using the same procedure as described above.

### Cell culture and fluorescence labelling

African green monkey kidney COS-7 cells (ATCC CCL-70) were cultivated in DMEM supplemented with 10% heat-inactivated FBS. Cells were maintained in a humidified incubator supplemented with 5% CO_2_. Cells were seeded on 8 well glass-bottom cell culture dishes (Cat. No. 80807, #1.5H(170*µm ±* 5*µm*) D 263 M Schott glass, sterilized, Ibidi) at a confluency of about 70% at least one day prior to imaging. Cells were washed with pre-warmed (37^*°*^C) PBS and the medium was exchanged to phenol-red free Leibovitz’s L-15 medium (Gibco, Thermo Fisher), which was heated to 37^*°*^C for live cell imaging.

For fluorescence labeling, cells were first chemically fixed via the following PFA-PEM fixation protocol: We prepared PEM consisting of 80 mM PIPES, 5 mM EGTA, and 2 mM MgCl_2_ at pH 6.8. We then prepared a fixation solution of 4% PFA, 4% sucrose in PEM. The cells were fixed for 10 min in the fixation solution, which was pre-heated to (37^*°*^C) and then rinsed 3 times in PBS. The cells were reduced for 10 min with 50 mM NH_4_Cl at room temperature and then washed 3 times in PBS. The cells were permeabilized for 10 min with BSA 3% + 0.25%Triton X-100 in PBS and then washed 3 times with PBS before labeling.

For fluorescence labeling we used Phalloidin Alexa Fluor 647 (A22287, Thermo Fisher, ex./em. 660 nm/680 nm) and followed the online protocol for preparing a stock solution. We dissolved the vial content in 150 *µl* anhydrous DMSO to yield a concentration of approximately 66 *µM*. We then diluted 0.5 *µl* of the stock solution in 200 *µl* PBS and added it to the fixed cells and incubated for 45 min at room temperature. After washing with PBS 3 times, the sample was mounted in PBS and directly imaged on the microscope. No blinking buffer was needed or used.

## Acknowledgment

We thank Pierre Jouchet for sharing the responsibility of running the cell culture. We thank Andrew E. S. Barentine, Pierre Jouchet and Ashwin Balaji for fruitful discussions and feedback on this work. We thank Pierre Jouchet for proofreading part of this work and useful comments. This project was supported in part by the U. S. National Institute of General Medical Sciences, Grant No. R35-GM118067.

## Author contributions

W.E.M. supervised the project; M.K. initiated and conceived the research; M.K. designed the optical layout and built the optical system; M.K. developed the algorithm and implemented the corresponding software for hardware control; M.K. implemented the electronic control with guidance from W.E.M.; M.K. performed the experiments, analyzed the data and prepared the figures. M.K. and W.E.M. wrote the manuscript, participated in the discussions and data interpretation.

## Code and data availability statements

The python code utilized in this study is available at Stanford Digital Repository at https://doi.org/10.25740/yr405qh5532. The source data files for the analysis and additional timelapse videos as extended data are also provided in the Stanford Digital Repository.

## References

[1] Rust, M. J., Bates, M. & Zhuang, X. Sub-diffraction-limit imaging by stochastic optical reconstruction microscopy (storm). Nature Methods 3, 793–796 (2006).

[2] Heilemann, M. et al. Subdiffraction-resolution fluorescence imaging with conventional fluorescent probes. Angewandte Chemie International Edition 47, 6172–6176 (2008).

[3] Möckl, L. & Moerner, W. E. Super-resolution microscopy with single molecules in biology and beyond–essentials, current trends, and future challenges. Journal of the American Chemical Society 142, 17828–17844 (2020). Doi: 10.1021/jacs.0c08178.

[4] Betzig, E. et al. Imaging intracellular fluorescent proteins at nanometer resolution. Science 313, 1642–1645 (2006).

[5] Klar, T. A., Jakobs, S., Dyba, M., Egner, A. & Hell, S. W. Fluorescence microscopy with diffraction resolution barrier broken by stimulated emission. Proceedings of the National Academy of Sciences 97, 8206–8210 (2000).

[6] Gustafsson, M. G. L. Surpassing the lateral resolution limit by a factor of two using structured illumination microscopy. Journal of Microscopy 198, 82–87 (2000).

[7] Sheppard, C. Super-resolution in confocal imaging. Optik - International Journal for Light and Electron Optics 80, 53 (1988).

[8] Pawley, J. B. Handbook of Biological Confocal Microscopy (Springer US, 2006).

[9] Müller, C. B. & Enderlein, J. Image scanning microscopy. Physical Review Letters 104, 198101 (2010).

[10] Sheppard, C. J. R., Mehta, S. B. & Heintzmann, R. Superresolution by image scanning microscopy using pixel reassignment. Optics Letters 38, 2889–2892 (2013).

[11] Castello, M. et al. A robust and versatile platform for image scanning microscopy enabling super-resolution flim. Nature Methods 16, 175–178 (2019).

[12] Zunino, A. et al. Structured detection for simultaneous super-resolution and optical sectioning in laser scanning microscopy. Nature Photonics 19, 888–897 (2025).

[13] York, A. G. et al. Instant super-resolution imaging in live cells and embryos via analog image processing. Nature Methods 10, 1122–1126 (2013).

[14] Roth, S., Sheppard, C. J. R., Wicker, K. & Heintzmann, R. Optical photon reassignment microscopy (opra). Optical Nanoscopy 2, 5 (2013).

[15] Luca, G. M. R. D. et al. Re-scan confocal microscopy: scanning twice for better resolution. Biomedical Optics Express 4, 2644–2656 (2013).

[16] Lindfors, K., Kalkbrenner, T., Stoller, P. & Sandoghdar, V. Detection and spectroscopy of gold nanoparticles using supercontinuum white light confocal microscopy. Physical Review Letters 93, 37401 (2004).

[17] Ginsberg, N. S., Hsieh, C.-L., Kukura, P., Piliarik, M. & Sandoghdar, V. Interferometric scattering microscopy. Nature Reviews Methods Primers 5, 23 (2025).

[18] Kukura, P. et al. High-speed nanoscopic tracking of the position and orientation of a single virus. Nature Methods 6, 923–927 (2009).

[19] Kashkanova, A. D., Blessing, M., Gemeinhardt, A., Soulat, D. & Sandoghdar, V. Precision size and refractive index analysis of weakly scattering nanoparticles in polydispersions. Nat. Methods 19, 586–593 (2022).

[20] Kashkanova, A. D., Albrecht, D., Küppers, M., Blessing, M. & Sandoghdar, V. Measuring concentration of nanoparticles in polydisperse mixtures using interferometric nanoparticle tracking analysis. ACS Nano 18, 19161–19168 (2024).

[21] Piliarik, M. & Sandoghdar, V. Direct optical sensing of single unlabelled proteins and super-resolution imaging of their binding sites. Nature Communications 5, 4495 (2014).

[22] Young, G. et al. Quantitative mass imaging of single biological macromolecules. Science 360, 423–427 (2018).

[23] Squires, A. H., Lavania, A. A., Dahlberg, P. D. & Moerner, W. E. Interferometric scattering enables fluorescence-free electrokinetic trapping of single nanoparticles in free solution. Nano Letters 19, 4112–4117 (2019).

[24] Delor, M., Weaver, H. L., Yu, Q. & Ginsberg, N. S. Imaging material functionality through three-dimensional nanoscale tracking of energy flow. Nature Materials 19, 56–62 (2020).

[25] Kueppers, M., Albrecht, D., Kashkanova, A. D., Luehr, J. & Sandoghdar, V. Confocal interferometric scattering microscopy reveals 3d nanoscopic structure and dynamics in live cells. Nature Communications 14, 1962 (2023).

[26] Shkarin, A. Alexshkarin/pylablib. Zenodo, https://github.com/SandoghdarLab/pyLabLib-cam-control (2023). Retrieved 2024-05-08.

[27] Linfoot, E. H. & Wolf, E. Phase distribution near focus in an aberration-free diffraction image. Proceedings of the Physical Society. Section B 69, 823 (1956).

[28] Cox, I. J., Sheppard, C. J. R. & Wilson, T. Improvement in resolution by nearly confocal microscopy. Applied Optics 21, 778–781 (1982).

[29] Duplinskiy, A., Frank, J., Bearne, K. & Lvovsky, A. I. Tsang’s resolution enhancement method for imaging with focused illumination. Light: Science & Applications 14, 159 (2025).

[30] Sheppard, C. J. R. & Choudhury, A. Image formation in the scanning microscope. Optica Acta: International Journal of Optics 24, 1051–1073 (1977).

[31] Novotny, L. & Hecht, B. Principles of Nano-Optics (Cambridge University Press, 2012), 2 edn.

[32] Roth, S., Sheppard, C. J. R. & Heintzmann, R. Superconcentration of light: circumventing the classical limit to achievable irradiance. Optics Letters 41, 2109–2112 (2016).

[33] Richards, B. & Wolf, E. Electromagnetic diffraction in optical systems, ii. structure of the image field in an aplanatic system. Proceedings of the Royal Society of London. Series A. Mathematical and Physical Sciences 253, 358–379 (1959).

[34] Kashkanova, A. D. et al. Precision single-particle localization using radial variance transform. Optics Express 29, 11070–11083 (2021).

[35] Zunino, A. et al. Open-source tools enable accessible and advanced image scanning microscopy data analysis. Nature Photonics 17, 457–458 (2023).

[36] Hsiao, Y.-T., Wu, T.-Y., Wu, B.-K.Chu, S.-W. & Hsieh, C.-L. Spinning disk interferometric scattering confocal microscopy captures millisecond timescale dynamics of living cells. Optics Express 30, 45233–45245 (2022).

[37] Zhitnitsky, A., Benjamin, E., Bitton, O. & Oron, D. Super-resolved coherent anti-stokes raman scattering microscopy by coherent image scanning. Nature Communications 15, 10073 (2024).

[38] Huff, J. The airyscan detector from zeiss: confocal imaging with improved signal-to-noise ratio and super-resolution. Nature Methods 12, i–ii (2015).

[39] Delattre, S. Igniting new confocal imaging potential – nikon ax r series with nsparc. Microscopy Today 31, 23–27 (2023).

[40] PicoQuant GmbH. Advancing flim with super-resolved spatial details and enhanced contrast: Novaism software for the luminosa microscope (2025). Accessed: 2025-09-18.

[41] Radmacher, N. et al. Doubling the resolution of fluorescence-lifetime single-molecule localization microscopy with image scanning microscopy. Nature Photonics 18, 1059–1066 (2024).

[42] Allan, D. et al. Trackpy v0.6.3. Zenodo, 10.5281/zenodo.12255 (2014). Retrieved 2025-09-18.

